# Smyd1 regulates H3K4me3 to support adaptive fatty acid oxidation in the adult heart during lipid overload

**DOI:** 10.64898/2026.11.18.763125

**Authors:** Risa Mukai, Shinichi Oka, Eun-Ah Sung, Koichiro Takayama, Soichiro Ikeda, Masato Matsushita, Takuma Takada, Junichi Sadohsima

**Author notes:** Correspondence to: Risa Mukai, Department of Cell Biology and Molecular Medicine, Cardiovascular Research Institute, Rutgers New Jersey Medical School, Newark, NJ, USA. Department of Cell Biology and Molecular Medicine, Cardiovascular Research Institute, Rutgers New Jersey Medical School, 185 South Orange Avenue, Newark, NJ 07103, USA.

## Abstract

Obesity and metabolic syndrome are major risk factors for heart failure, and impaired fatty acid oxidation (FAO) is a key contributor to metabolic dysfunction in obesity-related cardiomyopathy. However, how the heart maintains FAO and adapts to chronic lipid overload remains poorly understood.

Here, we identify SET and MYND domain-containing protein 1 (Smyd1), a muscle-specific histone methyltransferase, as an early compensatory regulator of cardiac metabolism in response to lipid overload. In wild-type mouse hearts, high-fat diet (HFD) feeding induced transient increases in Smyd1 expression and H3K4me3 levels during the early-to-mid phase of metabolic stress, both of which declined to baseline levels or below during prolonged HFD exposure. Single-cell RNA sequencing further revealed that this adaptive increase in Smyd1 expression occurred predominantly in cardiomyocytes. In contrast, cardiac-specific Smyd1 heterozygous knockout mice exhibited accelerated cardiac dysfunction, left ventricular dilation, increased myocardial fibrosis, and reduced survival following HFD feeding. Mechanistically, Smyd1 deficiency impaired the adaptive upregulation of key FAO-related genes and reduced cardiac FAO activity. Furthermore, co-immunoprecipitation analysis revealed an interaction between Smyd1 and peroxisome proliferator-activated receptor alpha (PPARα), suggesting a potential mechanism by which Smyd1 supports FAO gene expression.

Taken together, these findings identify Smyd1 as an early compensatory regulator that supports FAO-dependent metabolic adaptation and protects the heart against lipid overload-induced cardiac dysfunction. This adaptive mechanism, involving H3K4me3 regulation and the Smyd1–PPARα interaction, may provide potential therapeutic targets for therapeutic strategies aimed at preserving cardiac metabolic flexibility in obesity-related cardiomyopathy.

**Graphical Abstract:** 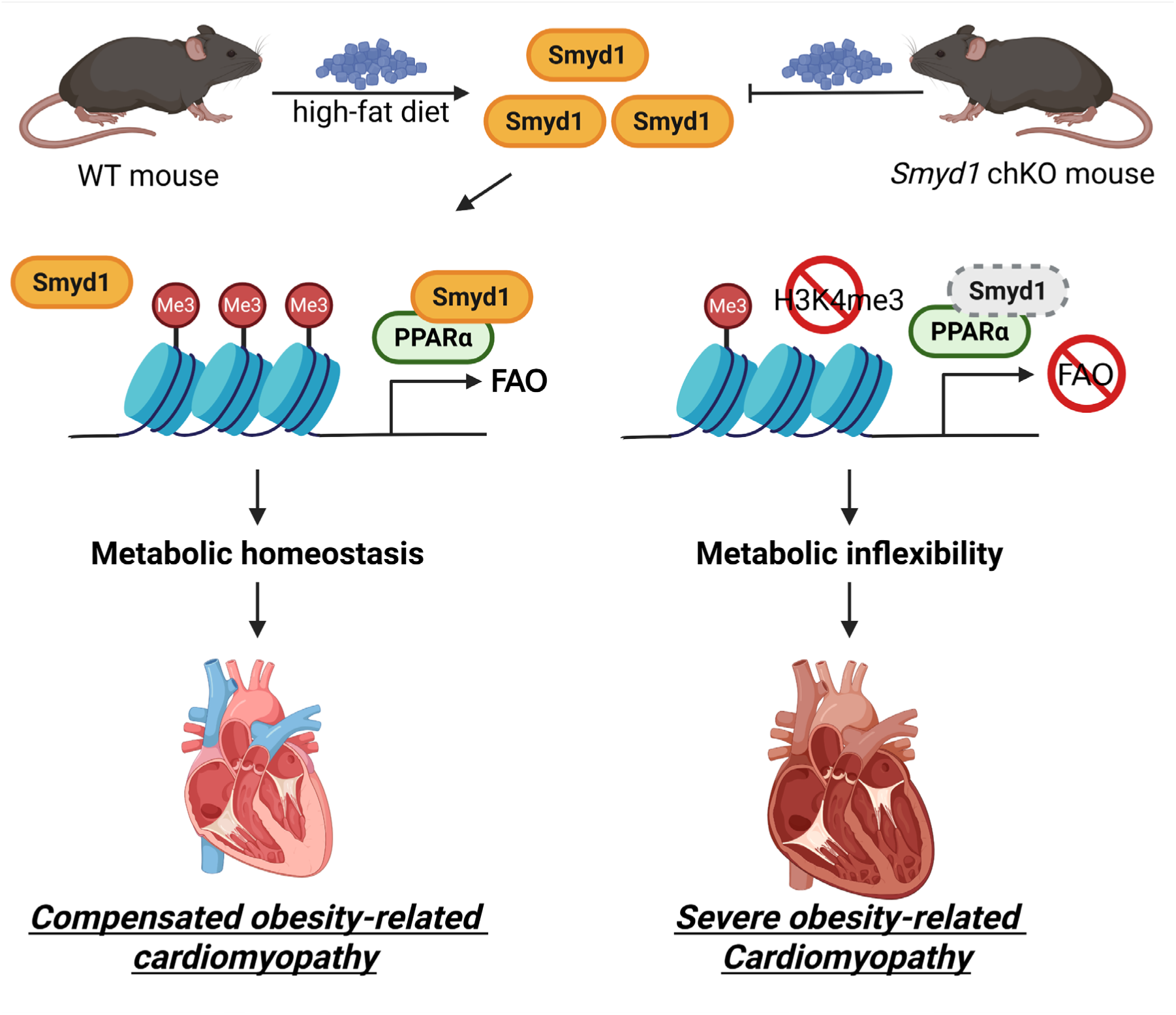

## 1. Introduction

The adult mammalian heart is a highly energy-demanding organ that relies on continuous ATP production to sustain contractile function^1^. Under physiological conditions, fatty acid oxidation (FAO) provides 60 to 90 percent of the cardiac ATP supply, with glucose and other substrates contributing to the remainder^1^. This balance between FAO and glucose oxidation is tightly regulated and allows the heart to adapt to varying energy demands^2^. However, in the setting of chronic nutrient excess, such as obesity and metabolic syndrome, this balance is disrupted, leading to altered substrate utilization, impaired mitochondrial efficiency, and eventual cardiac dysfunction^3^.

High-fat diet (HFD) feeding in rodents is a widely used model that mimics obesity-associated metabolic stress^4^. Prolonged HFD exposure leads to excessive lipid delivery to the myocardium, which initially triggers adaptive mechanisms aimed at increasing FAO capacity to match the higher lipid load^5^. However, when these adaptive responses fail or become insufficient, the resulting mismatch between lipid influx and oxidative capacity can cause toxic lipid accumulation, oxidative stress, and maladaptive remodeling^6^. Understanding the molecular regulators that control this early adaptive phase may reveal therapeutic targets for preventing the transition from compensated adaptation to decompensated heart failure.

Epigenetic regulation, particularly histone modifications, plays a central role in orchestrating transcriptional programs during metabolic stress^7^. Beyond mechanistic regulation, recent studies highlight that epigenetic marks, including DNA methylation and histone modifications, may serve as predictive biomarkers in cardiovascular disease and heart failure^8^. Trimethylation of histone H3 lysine 4 (H3K4me3) is a well-established marker of transcriptionally active promoters, facilitating recruitment of the transcriptional machinery and RNA polymerase II^9^. Several studies have demonstrated that H3K4me3 levels are dynamically regulated in response to metabolic cues, influencing the expression of metabolic enzymes and stress-response genes^10–12^. Despite this, the upstream regulators of H3K4me3 in the adult heart and their contribution to metabolic adaptation remain incompletely understood.

Smyd1 (SET and MYND domain-containing protein 1) is a cardiac-enriched histone methyltransferase known to catalyze H3K4 methylation^13, 14^. Smyd1 is essential during embryonic and early postnatal heart development, where it regulates sarcomere assembly, myofibril stability, and cardiomyocyte maturation^13^. Germline deletion of Smyd1 results in embryonic lethality due to severe structural heart defects, underscoring its critical developmental role. Cardiac-specific deletion of Smyd1 in adult mice has been shown to induce heart failure^15, 16^, suggesting that Smyd1 continues to play a critical role in maintaining cardiac function beyond development.

Smyd1 has also been shown to modulate proliferation of human pluripotent stem cell-derived cardiac progenitors through β-catenin and ERK signaling^17^, underscoring its broader role in cardiac growth and metabolic regulation. Notably, recent studies have reported that Smyd1 expression is inducible by metabolic or thermogenic stimuli in non-cardiac tissues, such as adipose tissue, raising the possibility that Smyd1 may act as a nutrient-responsive epigenetic regulator^18^. However, much less is known about the role of Smyd1 in the adult heart, particularly in the context of metabolic challenges such as HFD feeding. Emerging evidence indicates that Smyd1 participates in mitochondrial regulation and energy metabolism, including modulation of key transcriptional regulators of cardiac energetics, such as PGC-1α and PPARα^15, 16, 19^. Recent work has shown that Smyd1a protects against ischemic injury by regulating mitochondrial cristae remodeling and respiratory supercomplex formation^20^, suggesting that Smyd1 may also play adaptive roles under stress conditions. Given these findings, Smyd1 may act as a nutrient-sensitive epigenetic regulator that links histone modification to metabolic adaptation in the adult heart.

Given the central importance of FAO in the adult heart and the transcriptional activation of metabolic enzymes during early HFD exposure, we hypothesized that Smyd1 contributes to the adaptive response to lipid overload by promoting H3K4me3 enrichment at FAO-related gene loci. This hypothesis is supported by the known biochemical function of Smyd1 as a histone methyltransferase and by the potential for epigenetic enzymes to act as nutrient-sensitive regulators of gene expression.

To test this hypothesis, we examined the temporal expression of Smyd1 and H3K4me3 levels in the hearts of male and female mice during HFD consumption. We then assessed the functional and structural consequences of Smyd1 deficiency using a cardiomyocyte-specific heterozygous knockout (*Smyd1* chKO) model under both normal diet and HFD conditions. Finally, we investigated the impact of Smyd1 deficiency on the expression of key FAO enzymes to determine whether the loss of Smyd1 disrupts metabolic adaptation to lipid overload. Through these experiments, we aimed to define a previously unrecognized role for Smyd1 as an epigenetic regulator of cardiac metabolic flexibility.

## 2. Methods

### 2.1 Mouse models

Cardiac specific heterozygous *Smyd1* knockout mice were generated by the Genome Editing Shared Resource at Rutgers, The State University of New Jersey. Using CRISPR/Cas9-mediated incision within the targeted region of the *Smyd1* gene followed by homologous recombination, a loxP-flanked wild-type (WT) minigene cassette containing exons 2 and 3 of *Smyd1* was inserted into the endogenous locus. This design allowed expression of the WT minigene until Cre-mediated recombination occurred. To obtain cardiac-specific *Smyd1* knockout (*Smyd1* chKO) mice, the conditional allele was crossed with *Myh6-MerCreMer* transgenic mice, which express tamoxifen-inducible Cre recombinase specifically in cardiomyocytes. Upon tamoxifen administration, MerCreMer excised the loxP-flanked WT minigene cassette, thereby deleting exons 2 and 3 and generating a Smyd1-null allele specifically in cardiac tissue. All animal experiments were approved by the Institutional Animal Care and Use Committee of the New Jersey Medical School, Rutgers, The State University of New Jersey. The animal use in this study also conformed to the Guide for the Care and Use of Laboratory Animals published by the US National Institutes of Health (8th edition, 2011). To euthanize the mice for tissue harvest, the animals were deeply anesthetized with isoflurane and then euthanized by cervical dislocation. All animal experiments used both male and female mice. The investigators were blinded to the genotype groups during the experiments.

### 2.2 High fat diet feeding

A 60% fat diet (high fat diet, HFD) was purchased from Research Diets (#D12492) and used to induce cardiac hypertrophy in some experimental mice. A normal chow diet (10% fat, ND) was used as the control. Mice were placed on either the ND or HFD at an appropriate age and were maintained on their assigned diet for up to 3 months.

### 2.3 Tamoxifen feeding

Mice were fed tamoxifen-containing diet (400 mg per kg chow) ad libitum, starting at 3-4 months of age for one week, and then were switched to normal chow for an additional week. This tamoxifen induction period was then followed by the HFD or ND experiments. The tamoxifen-containing chow was purchased from ENVIGO (#TD. 130855).

### 2.4 Primary culture of neonatal rat ventricular cardiomyocytes

Neonatal rats were deeply anaesthetized with isoflurane (5%, once for 5–10 min) inhalation and euthanized by quickly removing their hearts. Primary cultures of ventricular cardiomyocytes were prepared from 1-day-old Charles River Laboratories (Crl)/Wistar Institute (WI) BR-Wistar rats (Harlan Laboratories, Somerville, NJ, USA) and maintained in culture as described previously^21^. A cardiomyocyte-rich fraction was obtained by centrifugation through a discontinuous Percoll gradient.

### 2.5 Echocardiography

Echocardiography was performed on cardiac specific heterozygous *Smyd1* knockout (*Smyd1* chKO) and littermate control mice at different time points, after 0 weeks to 12 weeks of HFD consumption. Under continuous isoflurane inhalation general anesthesia (0.75–1%), transthoracic echocardiography was conducted using a high-resolution Micro-Ultrasound system (Vevo 3100, FUJIFILM VisualSonics Inc., Toronto, Canada). Two-dimensional guided M-mode measurements of the left ventricle (LV) internal diameter were obtained from at least three beats and then averaged. LV end-diastolic dimension (LVEDD) was measured at the time of the apparent maximal LV diastolic dimension. LV ejection fraction (LVEF) was analyzed using Vevo LAB software (FUJIFILM VisualSonics, Inc.) to evaluate systolic function. All analyses were performed in a blinded manner regarding mouse genotype.

### 2.6 Immunoblotting analysis

Cardiac tissue homogenates were prepared from the LV apex. Cell lysates were prepared from primary cultures of rat ventricular myocytes. Both the homogenates and the cell lysates were prepared in an immunoprecipitation assay buffer containing 150 mM NaCl, 1% NP-40, 1% sodium deoxycholate, 0.1% sodium dodecyl sulfate, and 25 mM Tris, pH 7.6, and supplemented with a protease inhibitor cocktail (Sigma, #P8340), and a phosphatase inhibitor cocktail (Cell Signaling Technology (CST), #5870). The antibodies used include Smyd1 (Novus Biological, NBP2-20428), H3K4me3 (CST, #9751), Cpt1b (Sigma, #SAB5700834), MCAD (Sigma, #SAB5701546), VLCAD (Sigma, #HPA019006), c-Myc (Invitrogen, #MA1-980), Vinculin (Sigma, #V9131), α-tubulin (Sigma, #T6199), anti-mouse IgG HRP (CST, #7076), anti-rabbit IgG HRP (CST, #7074), anti-mouse IgG HRP TrueBlot ULTRA (eBioscience, #18-8817-33), and anti-rabbit IgG HRP TrueBlot ULTRA (eBioscience, #18-8816-33).

### 2.7 Histological analysis

The heart tissue was washed with PBS, fixed with 4% phosphate-buffered paraformaldehyde, embedded in paraffin and cut into 10-µm-thick sections. Serial sections of the heart were stained with wheat germ agglutinin (WGA) for analysis of cardiomyocyte cross-sectional area (CSA) and Picrosirius Red for analysis of myocardial fibrosis. CSA was quantified by tracing the outlines of 100–200 randomly selected cardiomyocytes from the LV with a clear nuclear image, using five distinct images per samples. Apoptosis was evaluated using the TUNEL assay. Nuclear density was determined by manually counting DAPI-stained nuclei in ten randomly selected fields per animal. TUNEL-positive nuclei were quantified by examining the entire section under the same magnification. All analyses were performed in a blinded manner using a fluorescence microscope (Eclipse E800, Nikon) and ImageJ software (NIH, Bethesda, MD).

### 2.8 Real-time quantitative polymerase chain reaction

Total RNA was extracted from mouse hearts using TRIzol (Invitrogen, #15596026). Total RNA was converted to cDNA using PrimeScript RT Master Mix (Takara, #RR036). Primer sequences are shown below: *Smyd1*-forward: GTGAAGAACGCAAGAGGCAGCT; *Smyd1*-reverse: CTCCTTCACCACTTCCTGAGAG; *Cpt1b*-forward: TTTGGGAACCACATCCGCCAA; *Cpt1b*-reverse: TTATGCCTGTGAGCTGGCCAC; *Mcad*-forward: GAAGCTGATGAGGGACGCCA; *Mcad*-reverse: GCTTGGAGCTTAGTTACACGAGG; *Rps18*-forward: AGTCCCTGCCCTTTGTACACA; *Rps18*-reverse: CGATCCGAGGGCCTCACTA. Data were normalized with the RT-PCR result for GAPDH. Quantitative RT-PCR was performed on the CFX 96 RealTime PCR Detection System (Bio-Rad). The Ct value determined using CFX Manager Software (version 2.0, Bio-Rad) for all samples was normalized to GAPDH, and the relative fold change was computed by the comparative Ct (ΔΔCt) method.

### 2.9 Single cell RNA sequencing

Mice were fed HFD for 4 weeks, while control mice were maintained on ND. Adult mouse heart cells were isolated from ND-fed (n = 3) and HFD-fed (n = 3) mice using previously published protocols^22^. Freshly isolated heart cells were pooled into ND and HFD groups and immediately fixed using Evercode Fixation v2 kits (Parse Biosciences), according to the manufacturer’s instructions. Fixed cells were processed into two sublibraries using the Evercode WT Mini v3 kit (Parse Biosciences). Barcoding and library generation were performed according to the manufacturer’s protocols. Barcoded libraries were sequenced on an Illumina platform and data were processed using the Parse Biosciences computational pipeline. Digital gene expression matrices were analyzed using Trailmaker, Parse’s cloud-based analysis platform. Standard preprocessing steps, including quality control, normalization, dimensionality reduction, clustering, and cell type annotation, were performed within the Trailmaker interface. Differential gene expression analysis and data visualization were conducted in Trailmaker using built-in statistical tools.

### 2.10 FAO activity assay

FAO activity was analyzed using the Fatty Acid Oxidation Assay Kit (E-141L, Biomedical Research Service Center, University at Buffalo, Atate University of New York). Briefly, heart tissues were subjected to homogenization with ice-cold cell lysis solution containing 150 mM NaCl, 1% NP-40, 1% sodium deoxycholate, 0.1% sodium dodecyl sulfate, and 25 mM Tris, pH 7.6. The sample was then centrifuged at14,000 rpm for 5 minutes, the supernatant was harvested, and the protein concentration was determined with the Quick Start Bradford Assay kit (BioRad, #5000201). The sample was mixed with FAO substrates diluted with the FAO assay solution and incubated at 37 °C for 45 minutes. The absorbance at 492 nm was measured.

### 2.11 Luciferase assay

Reporter and expression plasmids were transfected into neonatal rat ventricular cardiomyocytes (12-well plate) using LipofectAmine 2000 (Invitrogen #11668019). The amounts of expression vectors transfected into cells per well were 0.3 μg PPRE-luciferase reporter plasmids and 1 μl of 20 μM siRNA-scramble or siRNA-Smyd1. The PPRE-luciferase reporter plasmid has been described previously^23^. To mimic HFD–induced lipid overload, palmitic acid (PA, 250 μM) was added for 6 hours under plasmid transfection conditions. Cardiomyocytes were lysed with 100 μl Reporter lysis buffer after 5 days of transfection. The luminescence reaction was started by adding 5 μl lysate to 50 μl Reaction buffer, and luminescence was measured for 10 seconds using an OPTOCOMP I luminometer (MGM Instruments, Inc.). The luminescence was normalized by protein content measured using the Quick Start Bradford Protein Assay kit.

### 2.12 Statistical analysis

Statistical analyses were conducted using the GraphPad Prism 9.0 software program. Data are presented as line graphs expressing mean ± S.E.M for the indicated number of experiments or as dot-plots expressing mean ± S.E.M for the indicated number of mice. The difference in means between 2 columns or multiple columns was evaluated using the 2-tailed unpaired Student’s *t*-test (for 2 columns), one-way ANOVA with Tukey’s post-hoc test (for multiple comparisons), or one-way ANOVA with Dunnett’s post-hoc test (for multiple comparisons to control). The difference in means between more than 2 groups was evaluated using two-way ANOVA with Tukey’s post-hoc test. A *p* value of less than 0.05 was considered statistically significant. The reported significance values are always two-sided. Normal distribution of data was confirmed by the Shapiro-Wilk test. Non-normally distributed data were evaluated by the non-parametric Mann-Whitney test after confirming equality of rank variances.

## 3. Results

### 3.1 Smyd1 and H3K4me3 are transiently induced during HFD feeding

To determine whether high-fat diet (HFD) consumption alters the expression of Smyd1 in the adult mouse heart, we performed western blot analyses on left ventricular (LV) lysates from male and female wild-type (WT) mice fed HFD for 0, 2, 4, 8, or 12 weeks. Smyd1 expression increased after 4 weeks of HFD feeding, remained elevated through 8 weeks, and subsequently declined below baseline by 12 weeks in both sexes (**Figure 1A**). Given that Smyd1 functions as a histone methyltransferase catalyzing trimethylation of histone H3 lysine 4 (H3K4me3), we next examined H3K4me3 levels in the same samples. H3K4me3 showed a similar temporal pattern, increasing between 4 and 8 weeks and declining to baseline levels or below by 12 weeks, closely matching the changes in Smyd1 expression (**Figure 1A**). These data suggest that the transient increase in H3K4me3 during the early and middle stages of HFD consumption may be driven by increased Smyd1 activity, which may promote transcriptional activation of adaptive gene programs, whereas the decline at 12 weeks may reflect the loss of this adaptive response as metabolic stress continues.

**Figure 1.**
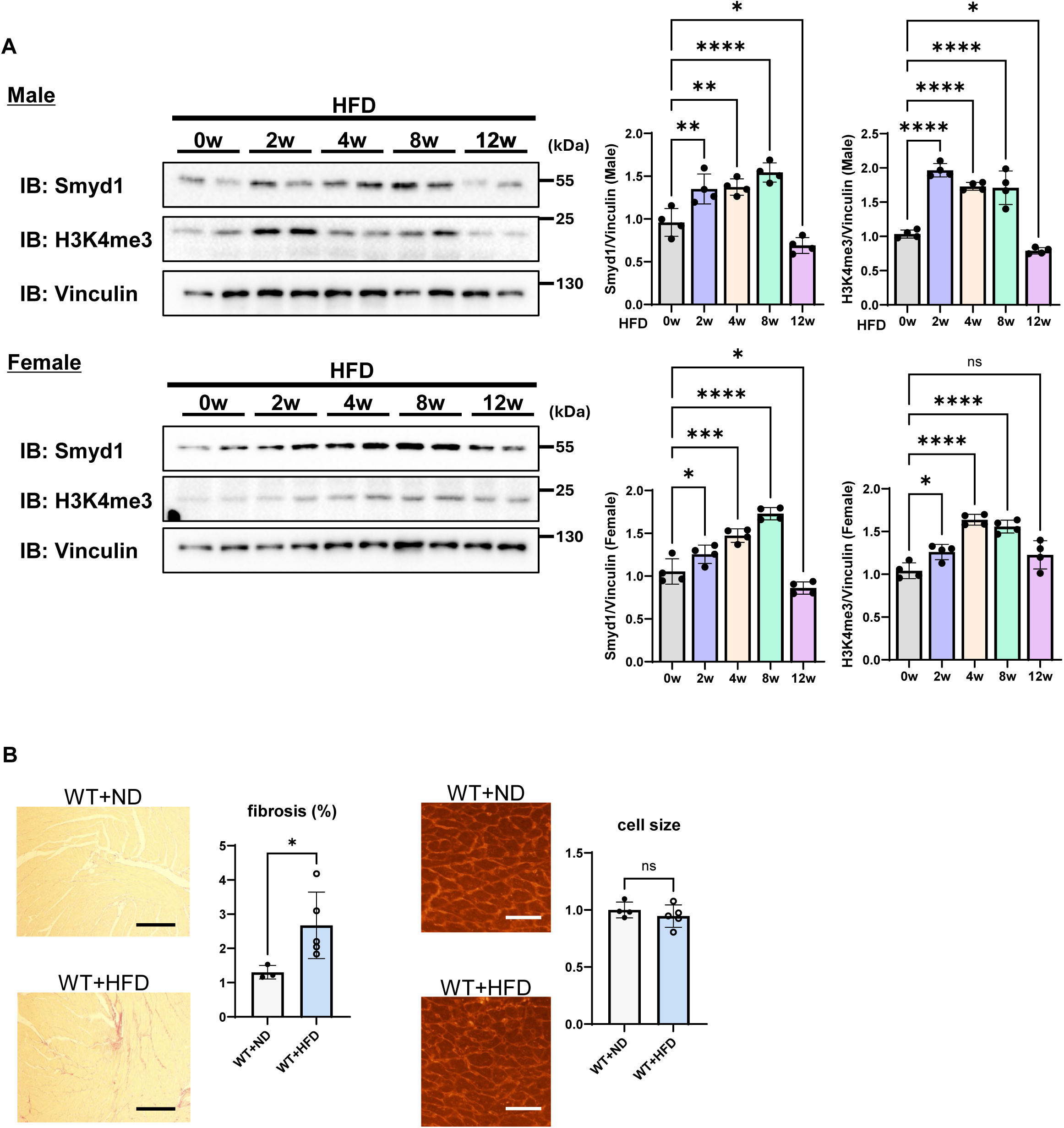
Time-dependent changes in Smyd1 expression and H3K4me3 levels in mouse hearts during HFD feeding. (**A**) Representative immunoblots showing H3K4me3 and Smyd1 protein expression in left ventricular tissue isolated from wild-type (WT) mice at 0, 2, 4, 8, and 12 weeks of HFD consumption. Vinculin was used as a loading control (n = 4). Data are presented as mean ± S.E.M. ns, not significant, \*\*\*\**p*<0.0001, \*\*\**p*<0.001, \**p*<0.05. Statistical significance was determined with one-way ANOVA with Dunnett’s post-hoc test. (**B**) Assessment of cardiac fibrosis and cardiomyocyte cell size. Representative histological images of picrosirius red–stained sections (left) and wheat germ agglutinin (WGA) staining of left ventricles (right) are shown, along with quantification of fibrotic area (% of total myocardial area) and cardiomyocyte size. Scale bar = 100 μm (left), 50 μm (right). Data are presented as mean ± S.E.M. ns, not significant, \*\*\*\**p*<0.0001, \*\*\**p*<0.001, \*\**p*<0.01, \**p*<0.05. Statistical significance was determined with one-way ANOVA with Dunnett’s post-hoc test (**A**) or 2-tailed unpaired Student’s *t*-test (**B**).

Echocardiographic assessment revealed no significant changes in LV ejection fraction (LVEF), LV diastolic dimension (LVDd) or interventricular septum thickness (IVSd) over the same time course (**Supplementary Figures S1A and S1B**). Necropsy data showed that the body weight/tibia length (BW/TL) ratio increased with HFD consumption, whereas heart weight/tibia length (HW/TL) and LV weight/tibia length (LVW/TL) remained unchanged (**Supplementary Figures S1C and S1D**).

Histological analyses were performed in WT mice to evaluate structural changes in the myocardium following 12 weeks of HFD feeding. Picrosirius red staining revealed an increase in interstitial collagen deposition in HFD-fed hearts, indicating enhanced cardiac fibrosis. In contrast, wheat germ agglutinin (WGA) staining showed no difference in cardiomyocyte cross-sectional area between control and HFD-fed groups, suggesting that HFD did not induce cardiomyocyte hypertrophy during this period. Taken together, these data indicate that 12 weeks of HFD primarily promotes fibrotic remodeling without causing measurable changes in cardiomyocyte size (**Figure 1B**).

### 3.2 Single-cell transcriptomic profiling reveals increased *Smyd1* expression during HFD consumption

To verify that a HFD induces Smyd1 upregulation in the heart, we performed single-cell RNA sequencing of mouse hearts after ND and HFD consumption **(Figure 2**). UMAP visualization revealed distinct clustering of cardiomyocytes under different diet conditions, with clear cluster shifts upon HFD consumption. Smyd1 upregulation was observed primarily in cardiomyocyte clusters that emerged under HFD conditions. In addition, dot plot analysis revealed increased Smyd1 transcript levels in HFD-fed mice compared with ND controls.

**Figure 2.**
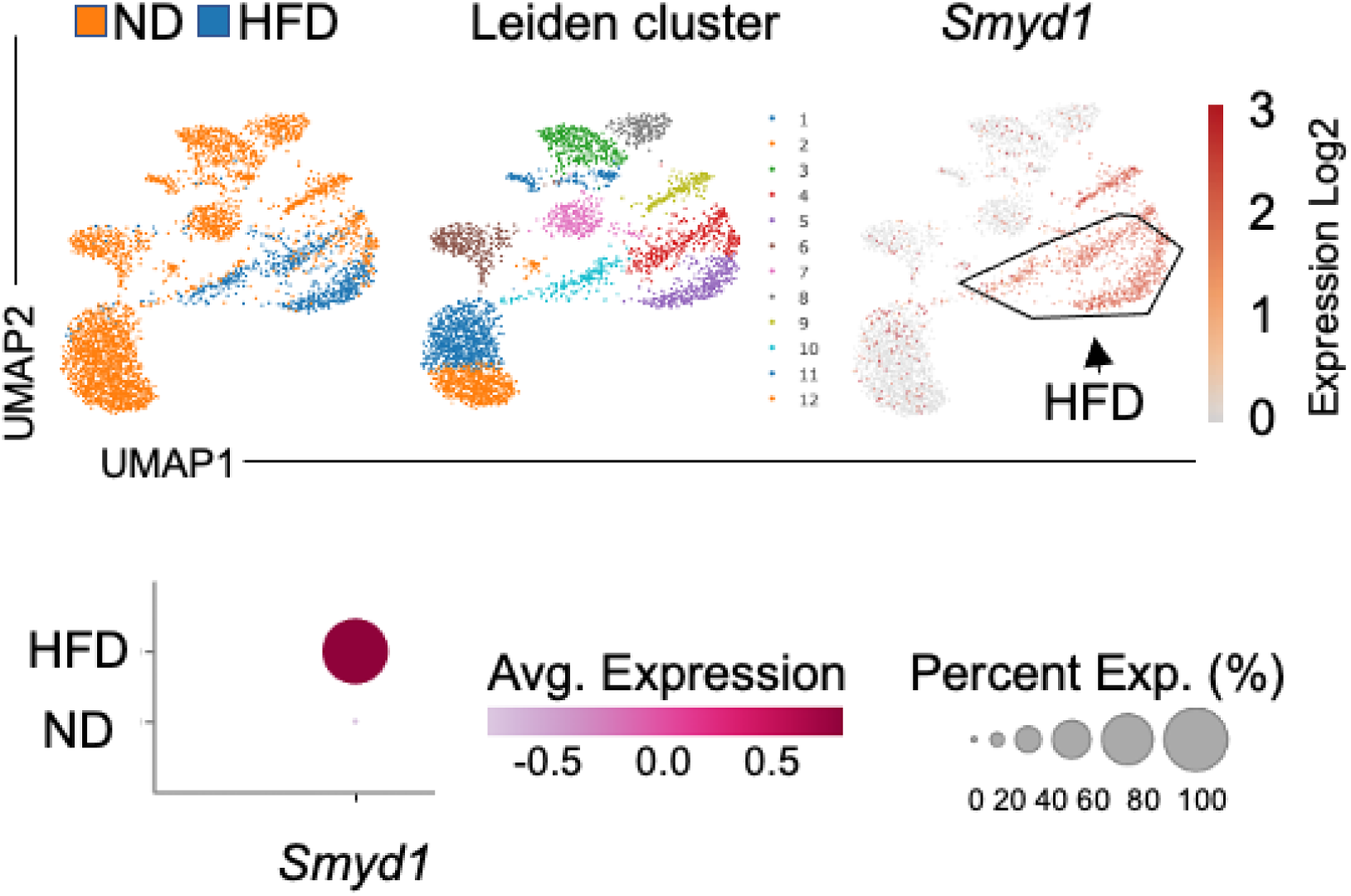
Single-cell RNA-seq analysis of Smyd1 expression in cardiomyocytes under ND and HFD conditions. UMAP plots of cardiomyocytes from ND- and HFD-fed mice, showing cluster annotations and the distribution of cells across diet conditions. Arrows indicate cluster shifts upon HFD exposure. UMAP plots show expression levels of *Smyd1* (upper panel). Dot plot visualization of *Symd1* from scRNA-seq data (bottom panel).

### 3.3 Loss of Smyd1 attenuates H3K4me3 induction and accelerates cardiac dysfunction during HFD consumption

To directly evaluate the functional role of Smyd1 in the heart, we generated cardiomyocyte-specific heterozygous *Smyd*1 knockout (*Smyd1* chKO) mice by crossing *Smyd1^flox/+^* mice with Myh6-MerCreMer transgenic mice and inducing recombination by tamoxifen feeding. Mice were then subjected to either the ND or HFD (**Figure 3A**). Western blot analysis confirmed a reduction of Smyd1 protein expression in *Smyd1* chKO hearts compared with *floxed* controls (*Smyd1^flox/+^*) (**Figure 3B**). Importantly, in *Smyd1^flox/+^* mice, HFD feeding induced a marked increase in both Smyd1 and H3K4me3 levels, but this HFD-induced rise in H3K4me3 was blunted in *Smyd1* chKO mice, indicating that Smyd1 is required for the increase in H3K4me3 during HFD consumption.

**Figure 3.**
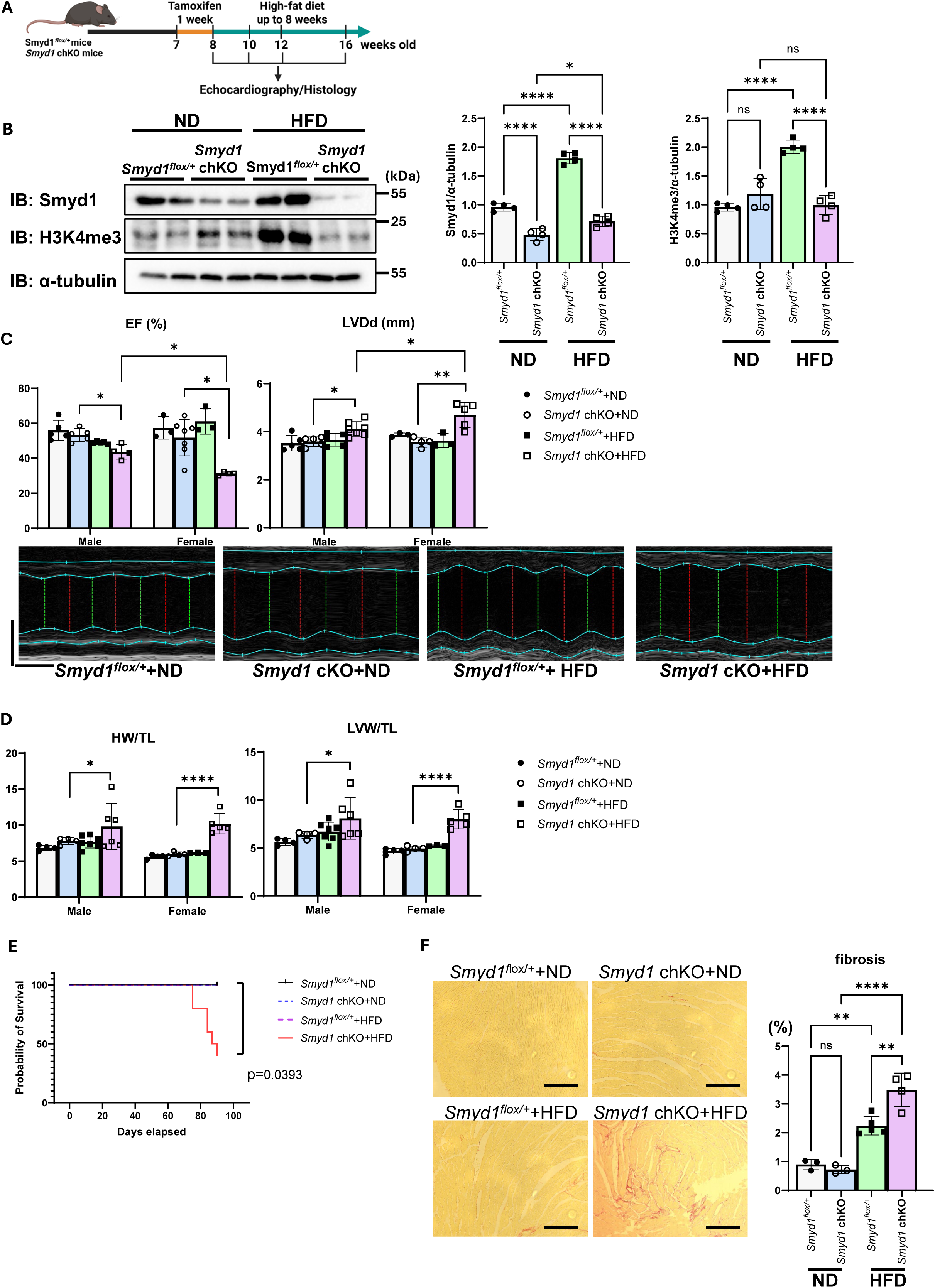
Cardiac-specific deletion of Smyd1 promotes HFD-induced cardiac remodeling. (**A**) Schematic representation of the experimental timeline. Both *Smyd1^flox/+^* (control) and *Smyd1* chKO mice were administered Tamoxifen for 1 week starting at 7 weeks of age to induce *Smyd1* gene deletion. Mice were then fed wither the ND or HFD for up to 8 weeks. (**B**) Representative immunoblot analysis of Smyd1 and H3K4me3 expression in hearts from *Smyd1^flox/+^* (control) and *Smyd1* chKO mice fed a normal diet (ND) or high-fat diet (HFD) for 8 weeks (n = 4). (**C**) Echocardiographic assessment of left ventricular ejection fraction (EF%) and left ventricular end-diastolic dimension (LVDd) in mice fed ND or HFD for 8 weeks (n = 3-7). Scale bar = 100 ms (horizontal), 2 mm (vertical). (**D**) Necropsy analysis in *Smyd1^flox/+^* and *Smyd1* chKO mice under ND or HFD conditions. Representative necropsy findings are shown (n = 3-7). (**E**) Kaplan–Meier survival curves of *Smyd1^flox/+^* and *Smyd1* chKO mice subjected to HFD feeding. Survival was significantly reduced in *Smyd1* chKO mice compared with controls (n = 11). (**F**) Assessment of cardiac fibrosis. Representative histological images of picrosirius red–stained sections of left ventricles are shown, together with quantification of fibrotic area (% of total myocardial area) (n = 3-5). Scale bar = 100 μm. Data are presented as mean ± S.E.M. ns, not significant, \*\*\*\**p*<0.0001, \*\**p*<0.01, \**p*<0.05. Statistical significance was determined with one-way ANOVA followed by Tukey’s post-hoc test (**B**, **C**, **D, F**), or two-way ANOVA with Tukey’s post-hoc test (**C**).

We next examined the impact of Smyd1 deficiency on cardiac function after 8 weeks of HFD consumption. Echocardiography revealed that LV ejection fraction (LVEF) was maintained in *Smyd1^flox/+^* mice during HFD consumption. In contrast, *Smyd1* chKO mice exhibited a significant decline in LVEF after HFD consumption (**Figure 3C**). Notably, the decline in LVEF during HFD consumption in *Smyd1* chKO mice was more severe in females than in males, suggesting a sex-dependent susceptibility to the loss of Smyd1 function during HFD consumption. LV diastolic dimension (LVDd) was not altered in response to HFD consumption in *Smyd1^flox/+^* mice. However, LVDd in *Smyd1* chKO mice was significantly greater after HFD consumption than after ND consumption. Furthermore, LVDd after HFD consumption was significantly greater in female than in male mice.

Consistent with the functional deterioration, necropsy analyses showed that both heart weight and LV weight were significantly increased in HFD-fed *Smyd1* chKO mice compared to in ND-fed counterparts in both sexes (**Figure 3D**). The mortality of mice was assessed by Kaplan–Meier survival curves. While *Smyd1^flox/+^* mice remained viable throughout the observation period, *Smyd1* chKO mice fed HFD showed a striking reduction in survival, with deaths beginning at around 2 months of HFD exposure. This indicates that the haploinsufficiency of Smyd1 accelerates the progression of heart failure during metabolic overload (**Figure 3E**). To evaluate the extent of cardiac remodeling, we performed picrosirius red staining of cardiac sections. *Smyd1^flox/+^* mouse hearts subjected to HFD displayed increased interstitial fibrosis compared with ND controls. Strikingly, *Smyd1* chKO hearts exhibited even greater fibrosis after HFD consumption (**Figure 3F**), suggesting that endogenous Smyd1 protects the heart against maladaptive remodeling.

### 3.4 Smyd1 deficiency abolishes HFD-induced upregulation of fatty acid oxidation enzymes

Since previous studies have suggested that Smyd1 regulates transcription of metabolic genes, including *PGC-1α*, *RXRα*, and *Perm1*^15, 24^, we next investigated the role of Smyd1 in regulating cardiac metabolism during HFD consumption. To examine whether Smyd1 directly regulates fatty acid oxidation (FAO) during metabolic stress, we assessed the expression and activity of key FAO enzymes in the heart of *Smyd1* chKO and control mice following ND and HFD consumption. Because FAO is the dominant energy source in the adult myocardium, adaptive upregulation of FAO enzymes is critical for maintaining energy homeostasis particularly when the lipid supply is increased and the use of glucose is limited in the presence of insulin resistance. We therefore hypothesized that loss of Smyd1 may blunt this metabolic adaptation. The qPCR analyses revealed that HFD consumption increases mRNA levels of *carnitine palmitoyltransferase 1b* (*Cpt1β*) and *medium-chain acyl-CoA dehydrogenase* (*Mcad*) in control hearts, but that this induction is abolished in *Smyd1* chKO mice (**Figure 4A**). Consistent with the mRNA results, western blot analyses demonstrated a robust upregulation of CPT1b, MCAD, and very long-chain acyl-CoA dehydrogenase (VLCAD) proteins in *Smyd1^flox/+^* mice, but not in *Smyd1* chKO mice (**Figure 4B**). Protein expression of these FAO enzymes was significantly lower in *Smyd1*-deficient hearts than in controls during HFD consumption, indicating that Smyd1 is required for the metabolic adaptation that enhances mitochondrial FAO capacity during HFD consumption.

**Figure 4.**
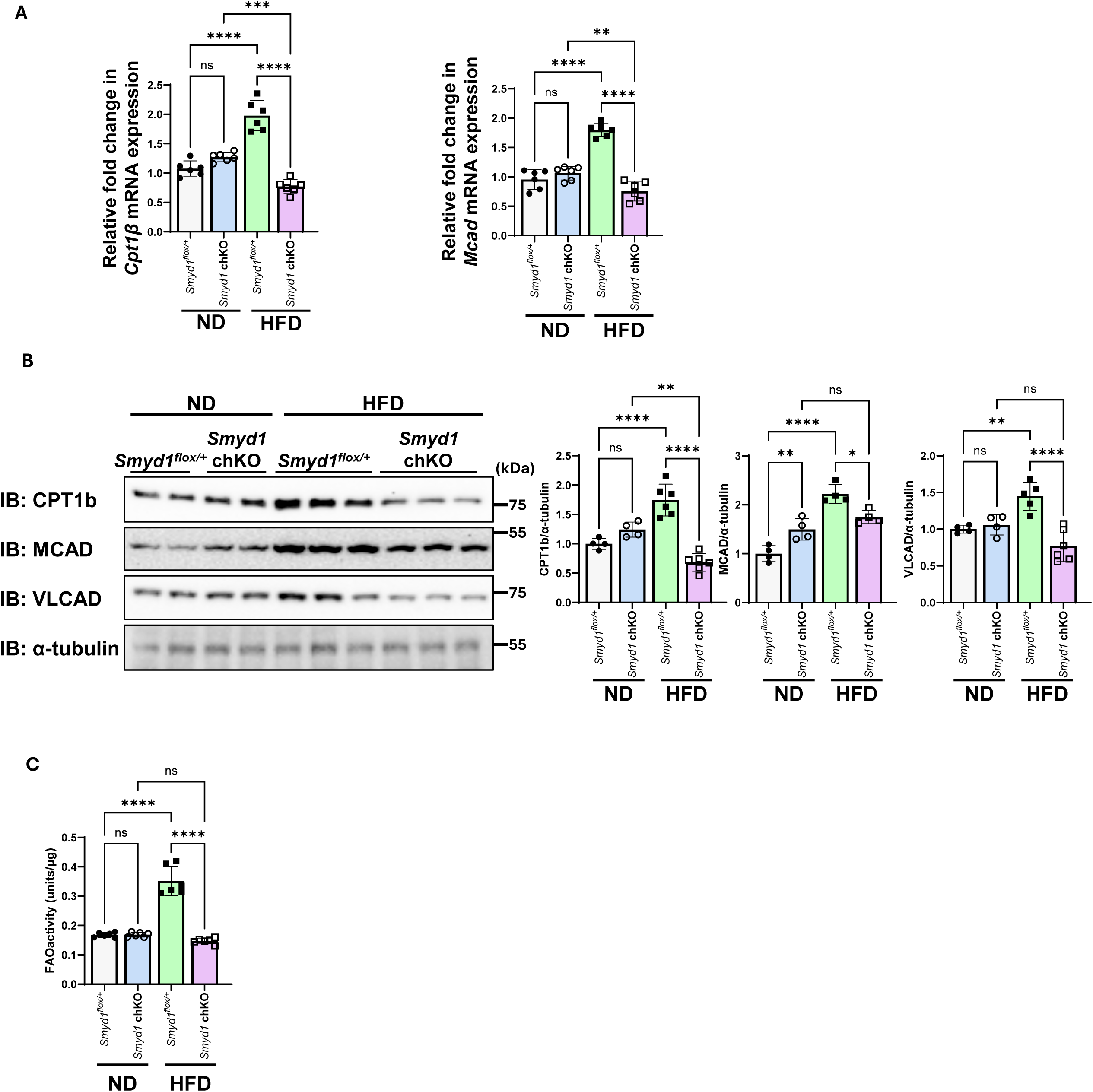
Impaired metabolic adaptation in Smyd1-deficient hearts. (**A**) Quantitative PCR analyses of *CPT1b* and *MCAD* mRNA expression in hearts from *Smyd1^flox/+^* and *Smyd1* chKO mice under ND and HFD conditions. Expression levels are normalized to control genes and presented as fold change (n = 6). (**B**) Representative immunoblots for CPT1b, MCAD, VLCAD in left ventricular lysates from *Smyd1^flox/+^* and *Smyd1* chKO mice fed a ND or HFD. α-tubulin was used as a loading control (n = 4). (**C**) Fatty acid oxidation (FAO) activity in left ventricular lysates from *Smyd1^flox/+^* and *Smyd1* chKO mice. FAO rates were measured using a commercially available FAO assay kit (n = 6). Data are presented as mean ± S.E.M. ns, not significant, \*\*\*\**p*<0.0001, \*\*\**p*<0.001 \*\**p*<0.01, \**p*<0.05. Statistical significant was determined with one-way ANOVA followed by Tukey’s post-hoc test (**A**, **B**, **C**).

To determine whether these molecular changes were reflected in metabolic function, we next evaluated FAO activity in cardiac lysates. A modest increase in FAO activity was detected in control hearts after HFD feeding, consistent with enhanced lipid utilization, but this response was attenuated in *Smyd1* chKO mice (**Figure 4C**), suggesting that Smyd1 deficiency compromises the overall FAO capacity in the heart.

### 3.5 Smyd1 physically interacts with PPARα and promotes PPARα transcription in response to palmitic acid

Because transcriptional activation of FAO genes is orchestrated by PPARα, we next investigated whether Smyd1 might physically associate with this nuclear receptor. Co-immunoprecipitation assays revealed a physical interaction between Smyd1 and PPARα (**Figure 5A**). This finding provides a mechanistic link between Smyd1-mediated epigenetic modification and PPARα-dependent transcriptional control of metabolic genes. It also suggests that Smyd1 may function as a transcriptional co-regulator that facilitates PPARα activity during metabolic stress.

**Figure 5.**
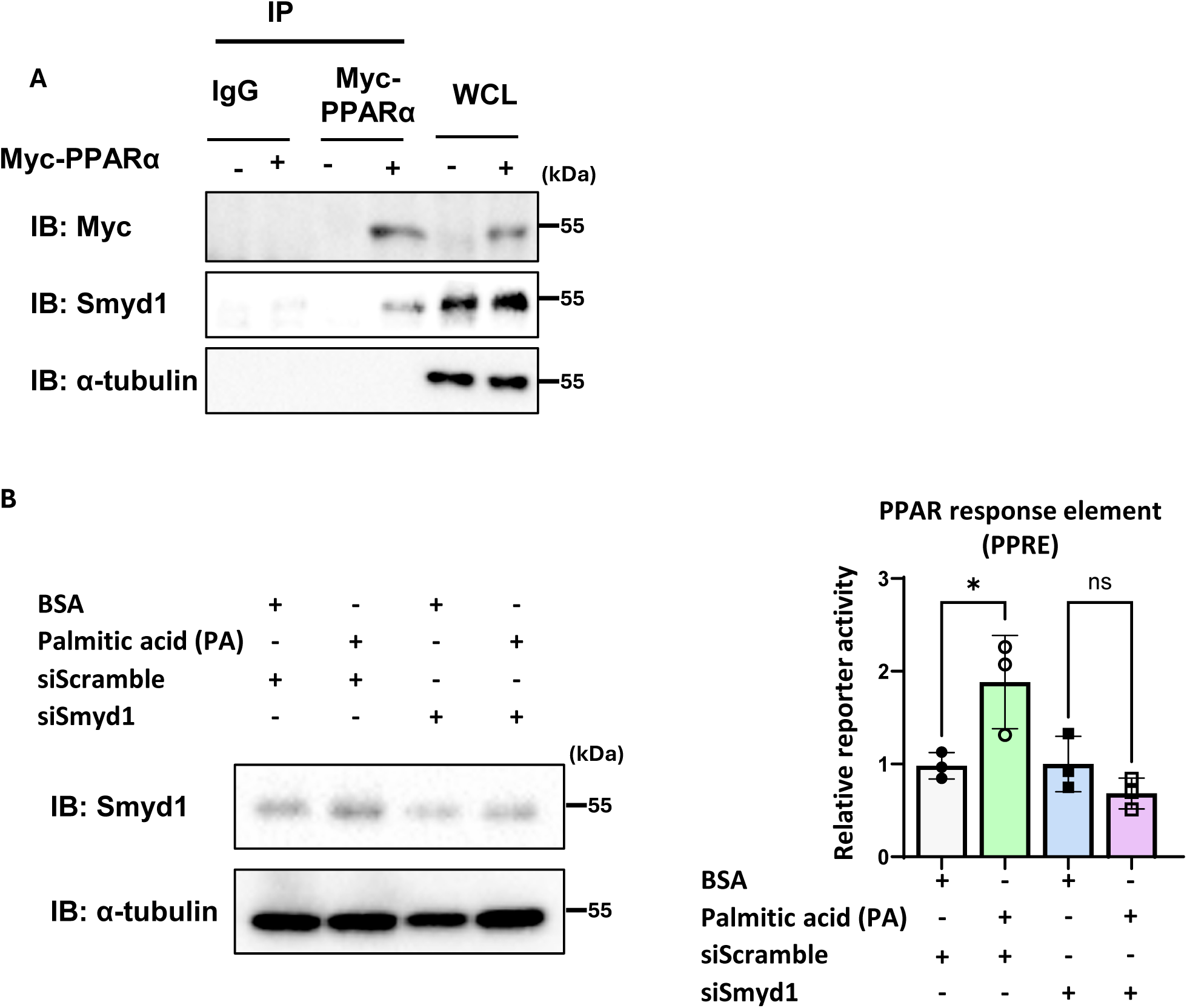
Smyd1interacts with PPARα to regulate palmitic acid–induced PPARα transcription. (**A**) Co-immunoprecipitation assays in mouse hearts using anti-Myc antibody. Detection of Myc-PPARα interaction with Smyd1. α-tubulin was used as a loading control (n = 3). (**B**) Cardiomyocytes were transfected with either siScramble or siSmyd1, followed by treatment with palmitic acid (PA) for 6 hours (n = 3). Data are presented as mean ± S.E.M. ns, not significant, \**p*<0.05. Statistical significance was determined with one-way ANOVA followed by Tukey’s post-hoc test (**B**).

To further determine whether Smyd1 modulates PPARα transcriptional activity, we performed luciferase reporter assays using a PPAR response element (PPRE)-luciferase reporter in neonatal rat cardiomyocytes. In the presence of palmitic acid (PA), which mimics lipid overload conditions, PPRE-luciferase activity was increased in control cells, consistent with activation of PPARα signaling. In contrast, this PA-induced activation was significantly reduced when Smyd1 was knocked down with siRNA, indicating that Smyd1 is required for PPARα transcriptional activity during metabolic stress (**Figure 5B**).

## 4. Discussion

Our data provide multiple lines of evidence that Smyd1 is an essential epigenetic regulator of FAO gene expression during HFD feeding. Smyd1 not only promotes H3K4me3 enrichment but also physically interacts with PPARα, forming a regulatory complex that enhances transcriptional activation of genes involved in FAO. A loss of Smyd1 disrupts this cooperative mechanism, resulting in diminished FAO enzyme expression, attenuated FAO activity, and impaired metabolic flexibility. These findings identify Smyd1 as a key molecular link between epigenetic regulation and transcriptional control of energy metabolism in the heart during nutritional stress.

In this study, we identify Smyd1 as a nutrient-responsive epigenetic regulator that supports metabolic adaptation in the adult heart during HFD consumption. We demonstrate that Smyd1 expression and the associated histone mark H3K4me3 are transiently induced in the myocardium during the early-to-mid phase of HFD exposure, coinciding with the period when the heart must adapt to increased lipid availability. This increase in Smyd1 and H3K4me3 was absent in *Smyd1* chKO mice, which exhibited impaired upregulation of key FAO enzymes, accelerated contractile dysfunction, increased fibrosis, and reduced survival, with females beings more severely affected than males. We observed the temporal dynamics Smyd1 and H3K4me3 peaking between 4 and 8 weeks of HFD consumption and returning to baseline by 12 weeks, suggesting that Smyd1-mediated chromatin remodeling may be part of a transient adaptive program. During the initial phase of HFD feeding, increased lipid delivery to the heart likely activates signaling pathways that induce Smyd1 expression, thereby enhancing H3K4me3 deposition at the promoters of FAO-related genes^25, 26^. This would facilitate increased transcription of *Cpt1β*, *Mcad*, and *Vlcad*, improving the heart’s capacity to oxidize fatty acids and maintain energy homeostasis. However, as HFD exposure persists, this adaptive program appears to wane, potentially due to chronic metabolic stress, epigenetic reprogramming toward a maladaptive state, or feedback inhibition from accumulated lipid intermediates^6^.

Functionally, our findings indicate that Smyd1 facilitates an adaptive transcriptional program that enhances fatty acid oxidation (FAO) capacity during lipid overload, thereby maintaining metabolic flexibility and protecting the heart from lipotoxic remodeling. Mechanistically, Smyd1 physically interacts with PPARα, a key transcriptional activator of FAO genes, providing a direct link between epigenetic regulation and nuclear receptor–mediated transcriptional control. Through this cooperation, Smyd1 likely promotes the recruitment of the transcription machinery to FAO promoters, enhancing transcriptional output during HFD consumption. In the absence of Smyd1, the myocardium fails to upregulate key FAO enzymes, leading to impaired fatty acid utilization, accumulation of lipid intermediates, and subsequent contractile dysfunction, promoting the transition from physiological adaptation to maladaptive heart failure. Although our current study reveals this functional interplay, the direct genomic targets of Smyd1 in the adult heart remain to be elucidated. Future ChIP-seq analyses will be essential to determine whether Smyd1 is recruited to FAO promoters or enhancers during metabolic stress. Moreover, it will be important to examine whether Smyd1 cooperates with other transcriptional regulators, such as ERRα, and whether it extends its regulatory influence beyond FAO to pathways controlling glucose and amino acid metabolism, thereby orchestrating a broader network of cardiac energy homeostasis.

Furthermore, our results indicate sex-specific differences, characterized by more pronounced dysfunction in female *Smyd1* chKO mice after HFD consumption, an intriguing observation that warrants further investigation^27^. Potential explanations include differences in baseline substrate utilization, hormonal modulation of mitochondrial function, and sex-specific transcriptional networks. Estrogen, for example, has been shown to enhance FAO and antioxidant capacity; thus, disruption of Smyd1-mediated transcription in females may disproportionately impair these protective pathways. Conversely, males may rely more heavily on other compensatory mechanisms that partially offset the loss of Smyd1.

An additional consideration is that the increased mortality observed in *Smyd1* chKO mice after 8 weeks of HFD feeding may be explained by the failure to sustain FAO upregulation. Our data show that Smyd1 expression returns to baseline levels after 8 weeks, which could contribute to the decline in FAO enzyme induction and the loss of metabolic flexibility at later stages of HFD exposure. This temporal relationship raises the possibility that Smyd1 is required not only for the initiation but also for the maintenance of FAO during prolonged metabolic stress. Once the adaptive FAO program collapses, persistent lipid overload likely promotes oxidative stress, mitochondrial dysfunction, and fibrosis, driving rapid progression to decompensated heart failure.

The parallel dynamics of Smyd1 expression and H3K4me3 deposition also suggest that Smyd1 partially governs this histone modification in the adult heart. While Smyd1 has been characterized as a histone methyltransferase, we cannot exclude the potential involvement of demethylase activity or other chromatin regulators. Future studies, including ChIP-seq and enzyme activity assays, are required to define the direct genomic targets of Smyd1 and the precise mechanisms by which it modulates chromatin landscapes under nutrient overload.

Clinically, the identification of Smyd1 as a regulator of cardiac metabolic flexibility has potential therapeutic implications. This is particularly relevant given recent metabolomic studies in HFpEF patients, which revealed altered lipid oxidation and amino acid metabolism as central features of human disease^28^. The impaired FAO adaptation we observed in Smyd1-deficient mice mirrors these clinical observations, suggesting translational relevance. Many patients with obesity or diabetes develop cardiomyopathy characterized by reduced FAO capacity and metabolic inflexibility. Pharmacological or gene therapy–based approaches that enhance Smyd1 activity, or mimic its downstream effects on chromatin accessibility, may offer new strategies to preserve cardiac metabolic resilience in the presence of obesity or diabetes. Conversely, understanding the conditions under which Smyd1 activity declines may provide insights into the transition from adaptive to maladaptive remodeling.

Endogenous Smyd1 levels are downregulated in the presence of heart failure, cardiac remodeling, and hypoxia. Under these conditions, metabolic adaptation to a HFD may be compromised, thereby facilitating the development of cardiomyopathy. If this is the case, molecular interventions aimed at restoring baseline Smyd1 expression could be considered. Although a recent study proposed the potential use of Smyd1 inhibitors for certain cardiovascular conditions^29^, such interventions should be approached with caution, as they may exert adverse effects in the context of metabolic stress.

In summary, our work reveals that Smyd1 acts as a nutrient-sensitive epigenetic regulator that transiently enhances FAO capacity during early HFD exposure. By maintaining efficient FAO and metabolic adaptability, Smyd1 preserves cardiac energetics and delays the onset of pathological remodeling. Loss of Smyd1 disrupts this adaptation, leading to impaired energy metabolism, accelerated pathological remodeling, and reduced survival. These findings highlight Smyd1 as a potential therapeutic target for preventing obesity-related cardiomyopathy and underscore the importance of epigenetic regulation in maintaining cardiac metabolic flexibility.

## Sources of Funding

This study was supported in part by U.S. Public Health Service grants HL91469, HL112330, HL138720, HL144626, and HL150881 (J.S.). This work was also supported by an American Heart Association Predoctoral Fellowship Award 915784 (E.-A.S.) and 25TPA 1481361 (J.S.), and by the Foundation Leducq Transatlantic Network of Excellence 15CVD04 (J.S.).

## Author contributions

R.M. and J.S. designed the experiments and wrote the paper; R.M. conducted the in vitro and animal experiments and analyses; S.O provided luciferase assay reagents and technical guidance; E.-A.S. isolated adult mouse cardiomyocytes and performed further single cell RNA-seq analysis; K.T., S.I., M.M, and T.T. conducted echocardiographic measurements of mice; R.M. performed histology analyses; J.S. generated project resources. All authors reviewed and commented on the manuscript.

## Acknowledgements

We thank Daniela Zablocki for critical reading of the manuscript.

## Conflict of interest

none declared

## Notes

### Competing Interest Statement

The authors have declared no competing interest.

