## Supplementary figures and images for "Smyd1 regulates H3K4me3 to support adaptive fatty acid oxidation in the adult heart during lipid overload"

### Supplement Figures

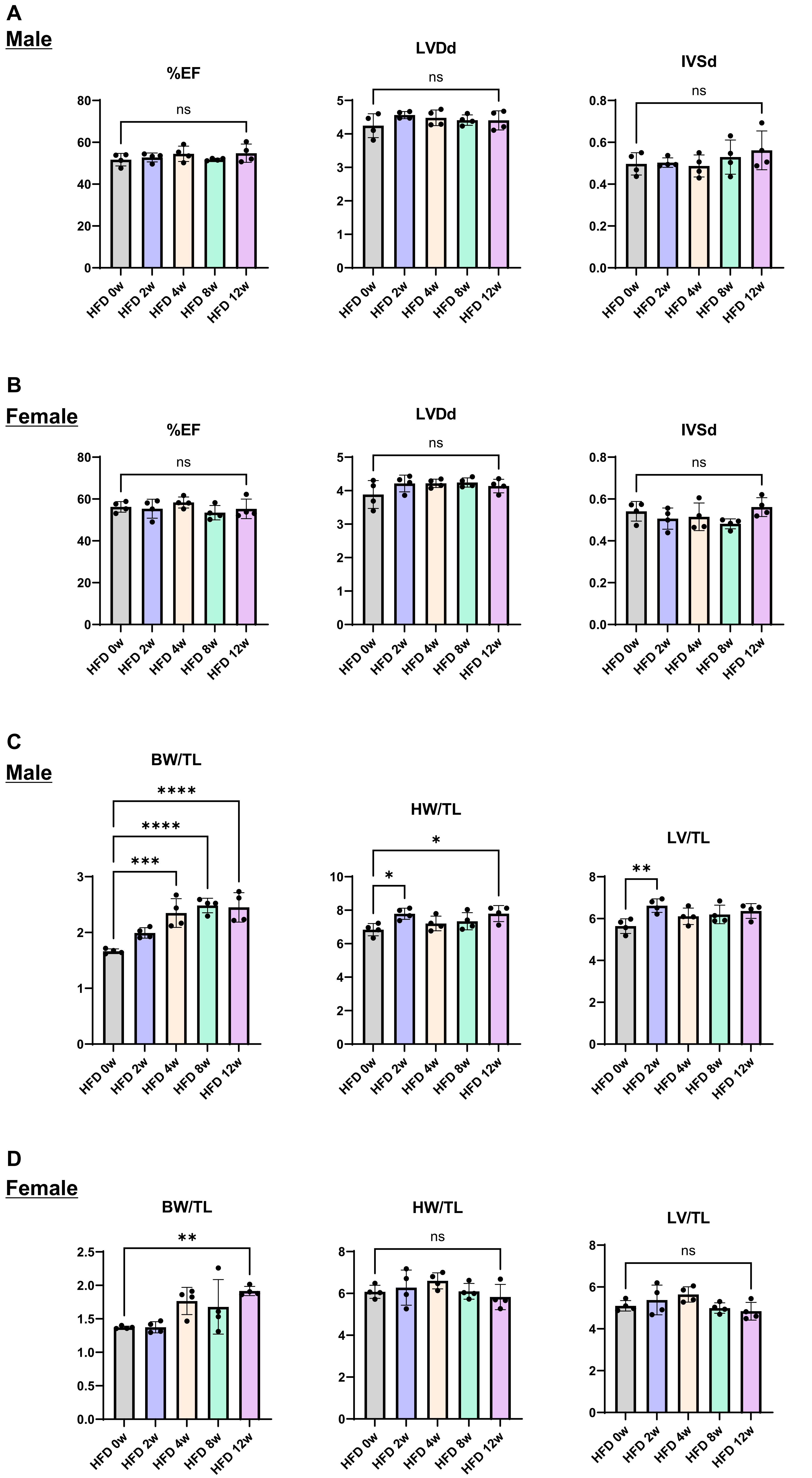


**Supplementary Figure 1**
